# *Efemp1* modulates elastic fiber formation and mechanics of the extrahepatic bile duct

**DOI:** 10.1101/2021.12.05.471313

**Authors:** Jessica Llewellyn, Emilia Roberts, Chengyang Liu, Ali Naji, Richard K. Assoian, Rebecca G. Wells

## Abstract

EGF-Containing Fibulin Extracellular Matrix Protein 1 (EFEMP1, also called fibulin 3) is an extracellular matrix protein linked in a genome-wide association study to biliary atresia, a fibro-inflammatory disease of the neonatal extrahepatic bile duct. EFEMP1 is expressed in most tissues and *Efemp1* null mice have decreased elastic fibers in visceral fascia; however, in contrast to other short fibulins (fibulins 4 and 5), EFEMP1 does not have a role in the development of large elastic fibers, and its overall function remains unclear. We demonstrated that EFEMP1 is expressed in the submucosa of both neonatal and adult mouse and human extrahepatic bile ducts and that, in adult *Efemp1^+/-^* mice, elastin organization into fibers is decreased. We used pressure myography, a technique developed to study the mechanics of the vasculature, to show that *Efemp1^+/-^* extrahepatic bile ducts are more compliant to luminal pressure, leading to increased circumferential stretch. We conclude that EFEMP1 has an important role in the formation of elastic fibers and mechanical properties of the extrahepatic bile duct. These data suggest that altered expression of EFEMP1 in the extrahepatic bile duct leads to an abnormal response to mechanical stress such as obstruction, potentially explaining the role of EFEMP1 in biliary atresia.

## Introduction

Biliary atresia is a rare and severe pediatric disease that initially causes fibrosis, obstruction, and obliteration of the extrahepatic bile ducts but rapidly leads to liver fibrosis and end-stage liver disease. The pathogenesis of biliary atresia is poorly understood. The inciting event appears to be exposure to an environmental agent such as a toxin ^[1, 2]^ or virus ^[3, 4]^ during gestation in a fetus with developmental and genetic susceptibility ^[5–7]^. Although no single gene has been found to be causative, multiple genes have been associated with biliary atresia in large population studies ^[8–16]^. One such gene is *EFEMP1*, which was identified in a genome-wide association study in two independent European-American biliary atresia patient cohorts as a candidate susceptibility gene ^[12]^.

EGF-Containing Fibulin Extracellular Matrix Protein 1 (EFEMP1), also known as fibulin-3, is member of the fibulin family of proteins. These proteins interact with a range of different matrix proteins, contributing to ECM organization and stability ^[17]^. Fibulin-3 is part of the subgroup of short fibulins that includes fibulins-4, −5 and −7, although its function is less well described. Fibulins-4 and −5 have critical roles in the formation of elastic fibers, particularly in large vessels. An elastic fiber is composed of an elastin core embedded in a fibrillin-rich microfibril scaffold. Its formation is a complex multistage process, with more than 30 elastic-fiber associated proteins identified ^[18]^. *Fibulin-4* deletions in mice cause perinatal lethality due to aortic rupture ^[19, 20]^. *Fibulin-5* deletion leads to irregular aggregation of elastin and lack of the normal characteristic lamellar structures in mouse aortas ^[21, 22]^. In contrast, fibulin-3 has a relatively low binding affinity for tropoelastin, the elastin monomer and, although fibulin 3 is expressed in lung and aorta, the large elastic fibers in these organs appear normal in *Efemp1* null mice ^[17, 23]^. *Efemp1* null mice instead demonstrate a loss of the fine elastic fibers found in fascia, with fascial herniation, early aging and low reproductive capacity, suggestive of a tissue- or elastic fiber size-specific function of EFEMP1. Here we establish a role for EFEMP1 in the formation of elastic fibers in the extrahepatic bile duct and demonstrate the mechanical importance of EFEMP1 in this tissue.

## Results

### EFEMP1 is highly expressed during the postnatal development of the extrahepatic bile duct in mouse and human

The expression pattern of EFEMP1 during the postnatal development of the mouse and human extrahepatic bile duct was assessed by immunostaining (Fig. 1A). In the neonatal mouse (P2) and human (3-day-old) extrahepatic bile duct, EFEMP1 is found highly expressed in both biliary epithelial cells and fibroblasts. In adult extrahepatic bile ducts however, EFEMP1 is mainly found deposited in the extrahepatic bile duct interstitium, with limited expression in cells. To evaluate a potential role for EFEMP1 in elastic fiber formation, we stained for both elastin and elastic fibers. Elastin deposition occurred later during postnatal development than EFEMP1 and was first apparent starting at P5, while elastic fibers only became visible between P10 and P14 (Fig. 1B). Interestingly although EFEMP1, elastin and elastic fibers are found throughout the extrahepatic bile duct submucosa, a strong deposition under the biliary epithelial layer is apparent in adult extrahepatic bile ducts (Fig. 1A-B). The sequential presence and similar distribution suggest a potential interaction between EFEMP1 and elastin.

**Figure 1:**
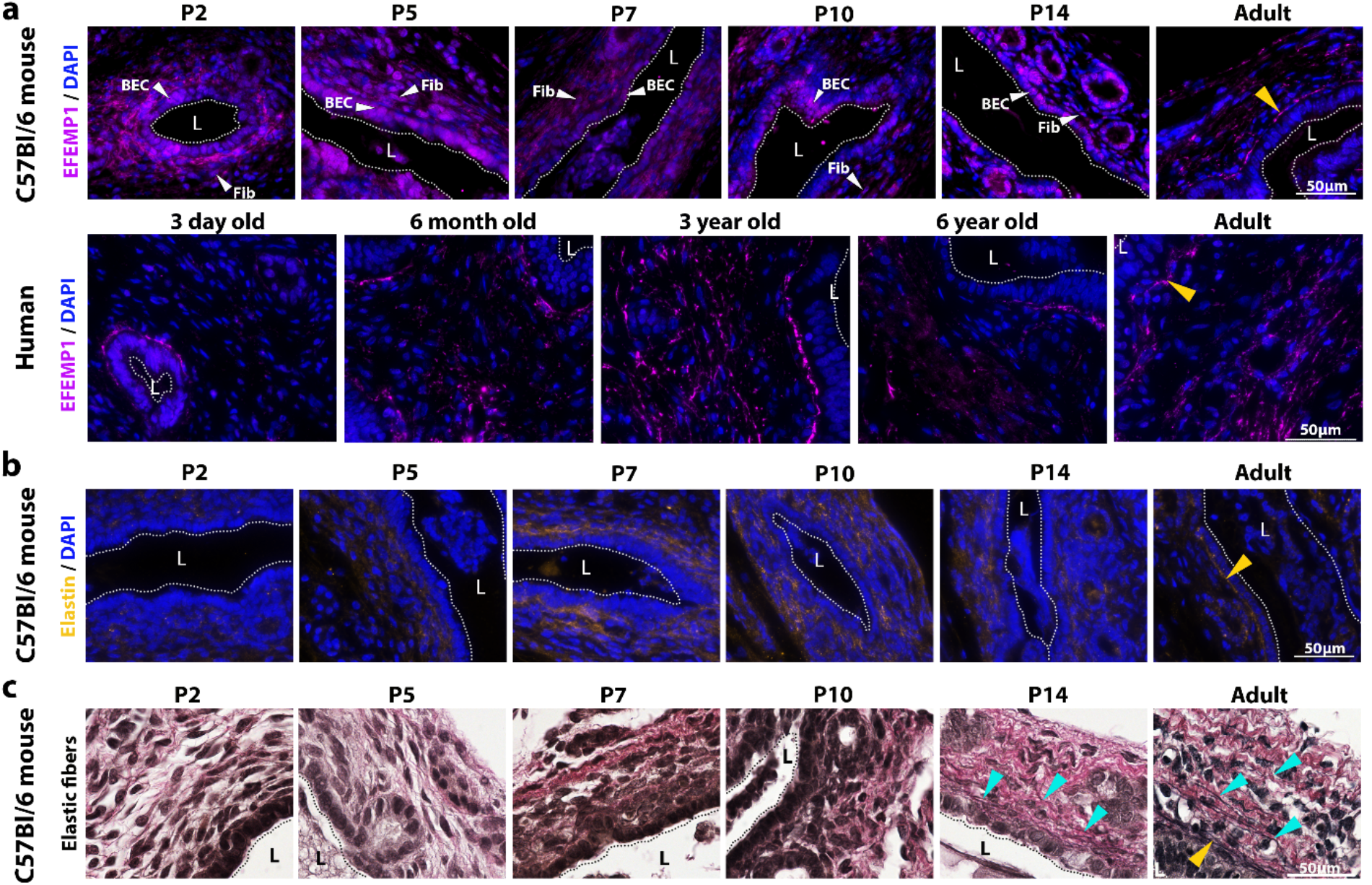
EFEMP1 expression in postnatal development of the mouse and human extrahepatic bile duct. (a) Representative images of EFEMP1, elastin and elastic fiber staining in neonatal (P2, 5, 7, 10, 14) and adult mouse extrahepatic bile duct as well as human extrahepatic bile duct from individuals aged 3 days old, 6 months old, 3 years old, 6 years old and 19 years old (adult). White arrows show biliary epithelial cells (BEC) and fibroblasts (fib) expressing EFEMP1. Yellow arrows show EFEMP1 localization under the biliary epithelial cell monolayer. (b) Representative images of elastin staining in mouse samples. In all immunohistochemistry staining, nuclei are stained with DAPI (blue). Yellow arrow shows elastin localization under the biliary epithelial cell monolayer (c) Representative images of elastic fiber staining, with stained elastic fibers in black (teal arrows), nuclei in dark brown and collagen in pink. Yellow arrow shows elastic fiber localization under the biliary epithelial cell monolayer. In all images, dotted lines and **L** denote lumens. For mice, 3-4 samples were stained for each age. Size bars = 50 μm.

### Efemp1^+/-^ extrahepatic bile ducts have reduced elastic fibers

To evaluate whether EFEMP1 has a role in elastic fiber formation in the extrahepatic bile duct, we compared expression of elastin and organized elastic fibers in the extrahepatic bile ducts of *Efemp1^+/-^* and WT mice. (Although *Efemp1* null mice were originally reported to be viable ^[23]^, *Efemp1^-/-^* mice were not born in our colony.) The amount of elastin deposition in the submucosa of the extrahepatic bile duct is comparable in the WT and *Efemp1^+/-^* mice, in both covering approximately 5% of the extrahepatic bile duct area; however, the distribution is different, with decreased localization under the biliary epithelial cell monolayer in the *Efemp1^+/-^* (Fig. 2A). The number of elastic fibers present in the extrahepatic bile duct submucosa is reduced by nearly half from 12.7 fibers/50 μm^2^ in the WT to 6.6 fibers/50 μm^2^ in the *Efemp1^+/-^* mice (Fig. 2B). This suggests that EFEMP1 has no impact on the expression and deposition of elastin but plays a role in the formation of elastic fibers. The expression of collagen, the other key structural protein in the extrahepatic bile duct, was examined using second harmonic generation (SHG) imaging, but both the amount of collagen and wavelength of collagen fiber crimps were similar in WT and *Efemp1^+/-^* extrahepatic bile ducts (Fig. 2C, Supplemental Fig. 1).

**Figure 2:**
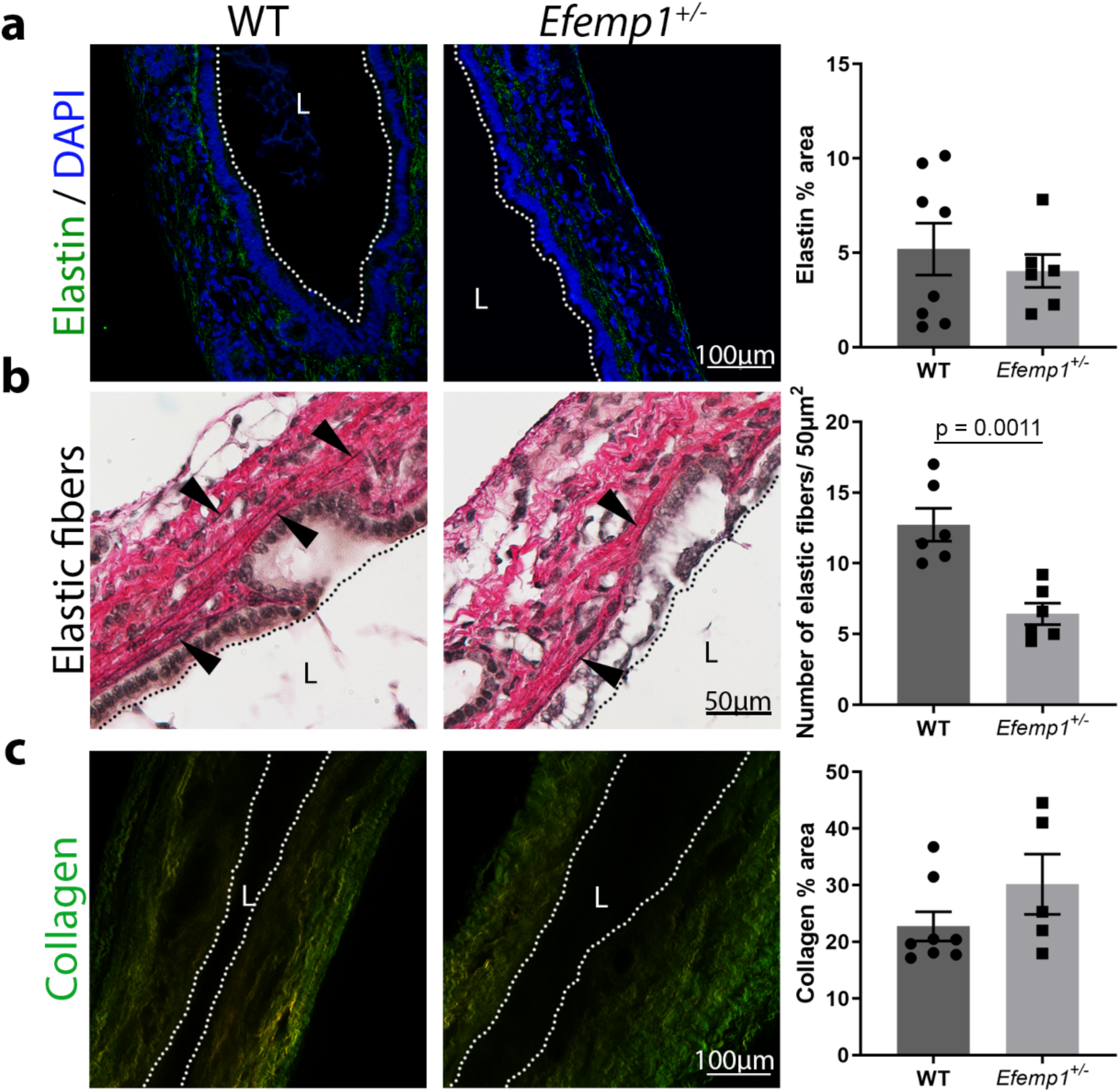
Elastic fibers but not elastin expression are reduced in adult *Efemp1^+/-^* extrahepatic bile ducts. (a) Representative staining of elastin in WT and *Efemp1^+/-^* extrahepatic bile ducts and quantification of percent area of staining. P= 0.5264 (b) Representative elastic fiber staining of WT and *Efemp1^+/-^* extrahepatic bile ducts and quantification of the number of elastic fibers per 50μm^2^. (c) Representative images of collagen by SHG imaging in WT and *Efemp1^+/-^* extrahepatic bile ducts, with quantification of percent area of signal. Forward signal in red and backward signal in green. P = 0.1843. Dotted lines and **L** denote lumens. 6-8 ducts were analyzed per genotype. Data shown are mean ± SEM, significance determined by unpaired T-test.

### Efemp1^+/-^ extrahepatic bile ducts are more compliant to luminal pressure

To determine the functional relevance of the changes in the extrahepatic bile duct submucosa observed with depletion of EFEMP1, we measured the mechanical properties of the extrahepatic bile duct using pressure myography. This technique, which was developed for and is typically used to study vessels, measures the changes in the diameter of a cylindrical structure in response to luminal pressure ^[24–27]^. When we compared the response of adult *Efemp1^+/-^* and WT extrahepatic bile ducts, we found that the *Efemp1^+/-^* ducts expanded more in response to pressure, leading to an increased outer diameter at any given pressure including at normal physiological pressures (0.8-3.5 mmHg) and at pressures found in obstructed ducts (12-15 mmHg) (Fig. 3A), using pressure values taken from the literature ^[28–31]^. Analysis was done on physiologically relevant pressures although data to 60 mmHg were collected (Fig. S2). The derived circumferential stretch-stress curve revealed a significant increase in stretch of the duct, while no statistically significant difference in stress was observed between WT and *Efemp1^+/-^* extrahepatic bile ducts (Fig. 3B). The stress-stretch curve has a similar shape as that of the aorta, in which the first phase is associated with elastin and the second with collagen ^[32, 33]^. The difference in stretch between WT and *Efemp1^+/-^* extrahepatic bile ducts occurs in the first low pressure phase as demonstrated by the 24.6% difference in luminal diameter between WT extrahepatic bile duct (75.5 ± 4.118 μm) and *Efemp1^+/-^* extrahepatic bile ducts (94.09 ± 8.239 μm) in the loaded (pressurized) state (Fig. 3C). The system is pressurized (unloaded to loaded) to near 0 mmHg (0.1 mmHg) at the start of each pressure myography test. The tangential modulus of the stress-stretch curve, an estimate of tissue stiffness, was also calculated, and the results indicated that stiffness of the extrahepatic bile duct significantly increases with increasing filling pressures. At physiologically relevant pressures, normal pressure (2 mmHg) and obstruction pressure (12 mmHg), the stiffnesses of WT and *Efemp1^+/-^* extrahepatic bile ducts are similar (Fig. 3D). These experiments demonstrate that EFEMP1 has a functional role in extrahepatic bile duct mechanics, in particular the circumferential stretch response to physiological filling pressures.

**Figure 3:**
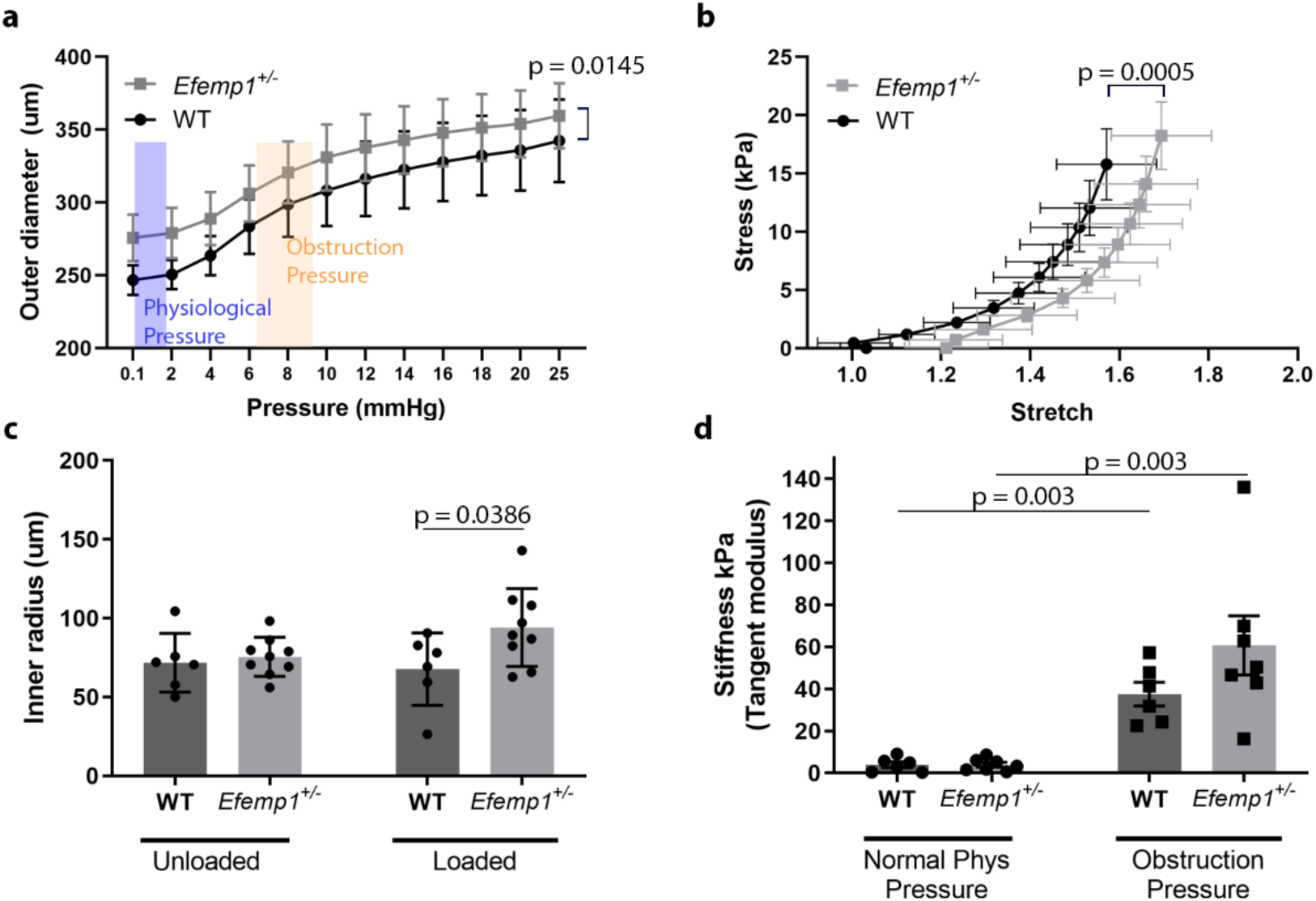
Mechanical properties measured by pressure myography are altered in *Efemp1^+/-^* extrahepatic bile ducts. (a) Measured outer diameter of extrahepatic bile ducts at increasing pressures. Physiological and obstruction pressures derived from the literature are indicated on the graph. (b) Calculated stress and stretch of WT and *Efemp1^+/-^* extrahepatic bile ducts under increasing pressure in 2-mm increments from 0-20 mm Hg. Significance shown is for stretch. No difference in stress was observed, p = 0.057. (c) Measured inner radius of unpressurized (unloaded) and pressurized (loaded; around 0.1 mmHg) WT and *Efemp1^+/-^* extrahepatic bile ducts. There is no difference between WT vs *Efemp^+/-^* in the unloaded state, p=0.9265 (d) Calculated tangential modulus/stiffness at normal physiological pressure (2 mmHg) and obstruction pressure (12 mmHg) in WT and *Efemp1^+/-^* extrahepatic bile ducts. No significant difference is observed comparing WT vs *Efemp1^+/-^* p = 0.5144 at both normal and obstruction pressures. 6-7 ducts were analyzed per genotype. Data shown are mean ± SEM, two-way ANOVA with a Šidák post-hoc test statistical test.

## Discussion

We have shown here that EFEMP1 is important for the formation of elastic fibers in the extrahepatic bile duct and contributes to duct mechanical properties, playing a significant role in the response to luminal pressure. These findings provide a potential mechanistic explanation for the link between *EFEMP1* and biliary atresia identified in a genome-wide association study ^[12]^, given that a controlled response to intraductal pressures and ability to resist obstruction are likely important in the function of the normal and injured duct.

Although it is likely to have a wide range of functions, EFEMP1 has been implicated in the formation and turnover of elastic fibers in a highly tissue-specific manner ^[34]^. The sequential expression of EFEMP1, elastin and then elastic fibers in postnatal development, the co-localization of the three under the biliary epithelial layer, and the reduction of the number of fibers in *Efemp1^+/-^* mice all suggest that EFEMP1 is important for the formation of elastic fibers in the extrahepatic bile duct. The lack of EFEMP1 expression in cells of the adult extrahepatic bile duct suggests that its role in elastic fiber homeostasis may be limited in the uninjured adult, although the mechanism by which EFEMP1 regulates elastic fibers requires additional investigation. EFEMP1 binds weakly to tropoelastin but does not bind fibrillin 1, the major component of the microfibril surrounding elastin ^[17, 35]^, and in the *Efemp1^-/-^* mouse, electron microscopy of the skin showed that the dark elastin core of elastic fibers was fragmented ^[23]^. This points towards a role for EFEMP1 in the formation of the elastin core of the elastic fiber.

To our knowledge, this study is the first to use pressure myography to test the mechanics of a non-vascular tissue. The extrahepatic biliary system is under very low pressure *in vivo*, particularly when compared to arteries, as reflected in the lack of thick elastin lamellae or a smooth muscle layer. To adapt the technique to the extrahepatic bile duct, lower pressures of 0-60 mmHg were used. Interestingly, while structure and pressures differ, the stretch experienced by the extrahepatic bile ducts is within the same range as experienced by arteries (stretch less than 2 fold) and both show a similar bi-phasic stress-strain curve. The initial phase has been attributed to the contribution of elastin and the second to collagen ^[32, 33]^. Given the similar degrees of stretch experienced by the extrahepatic bile duct and arteries, the two phases are likely to be similar for both structures. The curves for WT versus *Efemp1^+/-^* ducts show a difference in stretch which we attribute to changes in elastin; however, the second phases are remarkably similar, reflecting the similar collagen distribution and organization in the two genotypes.

Appropriate responses to mechanical stresses are key to maintaining tissue integrity and normal cell function. EFEMP1 has recently been linked to biliary atresia in a genome-wide association study. Our data show that even at physiological pressures the extrahepatic bile duct undergoes significant stretch, which we predict would affect both biliary epithelial cells and interstitial fibroblasts. Neonatal biliary epithelial cells in particularly have relatively immature tight junctions ^[36]^, and the increased stretch they experience in the setting of EFEMP1 abnormalities could result in the leakage of toxic bile into the submucosa, causing injury. Increased stretch could also impact submucosal fibroblasts, as fibroblasts in general are highly mechanosensitive cells and in tissues like skin and tendon show responses to stretch that include increased collagen 1 expression ^[37]^. Alternatively, the increased stretch observed with reduced EFEMP1 could serve a protective function, preventing the upstream (hepatic) consequences of obstruction.

In summary, EFEMP1 is involved in the formation of elastic fibers, which provide tensile strength limiting excessive mechanical disruptions to the tissue. An inability to respond appropriately to both physiological and obstruction pressures is likely to lead to cellular dysfunction and reduced ECM integrity. Although the predicted high binding promiscuity of EFEMP1 ^[34]^ suggests that it has as-yet-undescribed functions, the structural role of EFEMP1 in the extrahepatic bile duct may explain its link to biliary atresia.

## Experimental procedures

### Human tissue

Anonymized fresh human extrahepatic bile duct was obtained as part of the Human Pancreas Procurement and Analysis Program (HPPAP), which was granted IRB exemption (protocol 826489). Samples were taken after the unexpected deaths of otherwise healthy individuals and were formalin-fixed and embedded after receipt.

### Mouse tissue

All work with mice was in accordance with protocols approved by the University of Pennsylvania Institutional Animal Care and Use Committee, as per the National Institutes of Health Guide for the Use and Care of Animals. C57Bl/6j mice were obtained from the Jackson Laboratory (Bar Harbor, ME, USA). *Efemp1^+/-^* mice were a gift from Lihua Marmorstein (University of Arizona) ^[23]^. *Efemp1^+/-^* pairs were bred, however, although the nulls are reported to be viable, only *Efemp1^+/-^* and *Efemp1^+/+^* offspring were born in our colony and the heterozygotes were used for all experiments. Extrahepatic bile ducts were dissected from mice at different ages as noted in the figures and were formalin fixed (10%), paraffin embedded and sectioned at 5 μm thickness. Both male and female mice were used for all analyses.

### Immunofluorescence, elastic fiber staining and second harmonic generation imaging

For immunofluorescence of mouse and human extrahepatic bile ducts, samples were dewaxed with xylene and rehydrated through a graded series of alcohols and distilled water. Heat-mediated antigen retrieval with 10 mM citric acid pH 6.0 in a pressure cooker for 2 hrs was used for staining with antibodies against EFEMP1 (1:100, Thermo Fisher, PA5-29347, Waltham, MA); 0.5% hyaluronidase (Sigma, H3506 type I-S, St-Louis, MO) treatment for 60 min at 37°C was used for staining with antibodies against elastin (1:100, Cedarlane, CL55041AP, Burlington, NC). Sections were blocked with StartingBlock™ T20/phosphate buffered saline (PBS) Blocking Buffer (Thermo Scientific, Waltham, MA) before being incubated with primary antibodies (in 0.2% Triton X-100, 3% serum, in PBS) at 4°C overnight. Sections were then incubated with Cy3-conjugated anti-rabbit secondary antibody for 60 min (1:500, Vector Laboratories, Burlington, CA), DAPI for 10 min (4’, 6-Diamidino-2-Phenylindole, Dihydrochloride) and were mounted. Stained sections were imaged using a Zeiss Axio Observer 7 inverted microscope and ZEN blue software. For elastic fiber staining, paraffin sections were dewaxed and rehydrated and stained using an Elastic Stain Kit (Millipore, HT25A, Burlington, MA). Brightfield imaging was done on a Nikon E600 microscope with Nikon NIS-Elements software. For SHG imaging, paraffin sections were imaged using a Leica SP8 confocal/multiphoton microscope and Coherent Chameleon Vision II Ti:Sapphire laser (Leica, Buffalo Grove, IL) tuned to a wavelength of 910 nm.

### Image analysis

Raw image files from immunofluorescence and SHG imaging were processed using Fiji ImageJ software ^[38]^. Quantification of the percent area stained was assessed using the threshold and percent area functions. The number of elastic fibers per 50 μm^2^ was counted manually using the Count tool. The crimp wavelength and depth were measured manually using the Line tool. For all analyses, 3-5 images per sample were selected. Analyses were done blinded.

### Pressure myography

Extrahepatic bile ducts were isolated from adult (10-15 week old) WT and *Efemp1^+/-^* mice and placed in Hank’s balanced salt solution (HBSS) without calcium and magnesium at 37°C. The extrahepatic bile ducts were then carefully placed onto two 385 μm stainless steel cannulas on a DMT model 114P Pressure Myograph (DMT-USA, Ann Arbor, MI) and tied into place with 6-0 silk sutures (SUT-S 104, Braintree Scientific Inc., Braintree, MA), creating a closed system. The mounted ducts were stretched until straight, the force transducer was brought to zero, and then the ducts were stretched axially to 1 mN of force to standardize baseline axial conditions. The system was pressurized using medical air (AI USP 300; Airgas USA, Cherry Hill, NJ) and the extrahepatic bile ducts were preconditioned for 15 min at 40 mmHg before being pressurized in a stepwise manner from 0-20 mmHg in 2 mmHg increments and then to 60 mmHg in 5 mmHg increments. The system was fitted with a Nikon Diaphot inverted microscope, enabling measurement of the vessel length (VL), wall thickness (T) and outer diameter (OD) using MYOVIEW software (DMT-USA, Ann Arbor, MI). This was done 3 times on each extrahepatic bile duct and values were averaged for data analysis. No difference in initial diameter of pressurized extrahepatic bile ducts (0.1 mmHg) was observed amongst tests for a given duct, suggesting that duct returns to normal after each pressurization. Axial stretch measurements were not performed as these are unlikely to be forces experienced by the extrahepatic bile duct due to the low pressures of the system. Real-time imaging and the MYOVIEW software were used for direct measurement of the loaded outer diameters (*2a_o_*). From this, the unloaded inner radius (*A_i_*) was calculated as *A_i_*=*A_0_-H*, where H is the unloaded wall thickness. The duct volume (*V*) was used to determine the loaded inner radii using 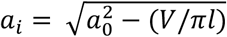 where l is the length between the inner sutures and measured using calipers. The loaded wall thickness (*h*) calculated using *h*=*a_0_*-*a_i_*. The circumferential stretch (λ_θ_) was calculated using λ_θ_= (*a_i_* + *h/2*)/(*A_i_*+*H/2*) and circumferential stress using *σ_ϑ_*= (*Pa_i_*)/*h* where P is pressure. As the extrahepatic bile duct shows a nonlinear stress-stretch relationship, the tangent moduli (slope) were calculated as a measure of extrahepatic bile duct stiffness. An exponential curve was fitted to the stress-stretch curve using MATLAB (R2021a, curve fitting tool). The tangent modulus was obtained by taking the derivative of the stress-stretch curves at normal physiological (2 mmHg) and obstruction (12 mmHg) pressures.

### Statistical analysis

Statistical significance was calculated with Prism 9 (GraphPad software, LaJolla, CA). The image analysis data were assessed by unpaired T-test. Statistical significance of the pressure myography was assessed by two-way ANOVA and Šidák post hoc analysis. Significant (p<0.05) results are noted on figures.

## Author Contributions

J.L. contributed to study concept and design, data acquisition, analysis and interpretation, and drafting of the manuscript. E.R. contributed to acquisition, analysis, and interpretation of data. A.N. and C.L provided material support and contributed to data interpretation. R.K.A contributed to study concept and design and participated in data interpretation. R.G.W. conceived ideas and designed the research, provided critical revision of the manuscript, obtained funding and supervised the study.

## Funding and Acknowledgments

We thank Lihua Y. Marmostein for the gift of *Efemp1^+/-^* mice. We gratefully acknowledge the assistance of the following core facilities and individuals at the University of Pennsylvania: the Penn Vet Imaging Core (NIH grant S10 OD021633-01) and Gordon Ruthel, the Perelman School of Medicine Cell and Developmental Biology Microscopy Core, and the NIDDK Center for Molecular Studies in Digestive and Liver Diseases Molecular Pathology and Imaging Core (NIH P30 DK050306). E.R. was supported by the Program in Translational Biomechanics of the Institute for Translational Medicine and Therapeutics at the University of Pennsylvania. Funding was provided by NIH R01 DK119290 (to R.G.W.); NIH grant UC4 DK112217 (to A.N.) and the NIDDK/City of Hope Integrated Islet Distribution Program (IIDP) Inst. #10039645; the Center for Engineering MechanoBiology (CEMB), an NSF Science and Technology Center, under grant agreement CMMI: 15-48571 (to R.G.W. and R.K.A.); and the Fred and Suzanne Biesecker Pediatric Liver Center (to R.G.W.).

## Abbreviations

EFEMP1: EGF-Containing Fibulin Extracellular Matrix Protein 1
SHG: second harmonic generation
DAPI: 4’,6-diamidino-2-phenylindole
WT: wild type
PBS: phosphate buffered saline
SEM: standard error of the mean

---

**Supplemental Figure 1:**
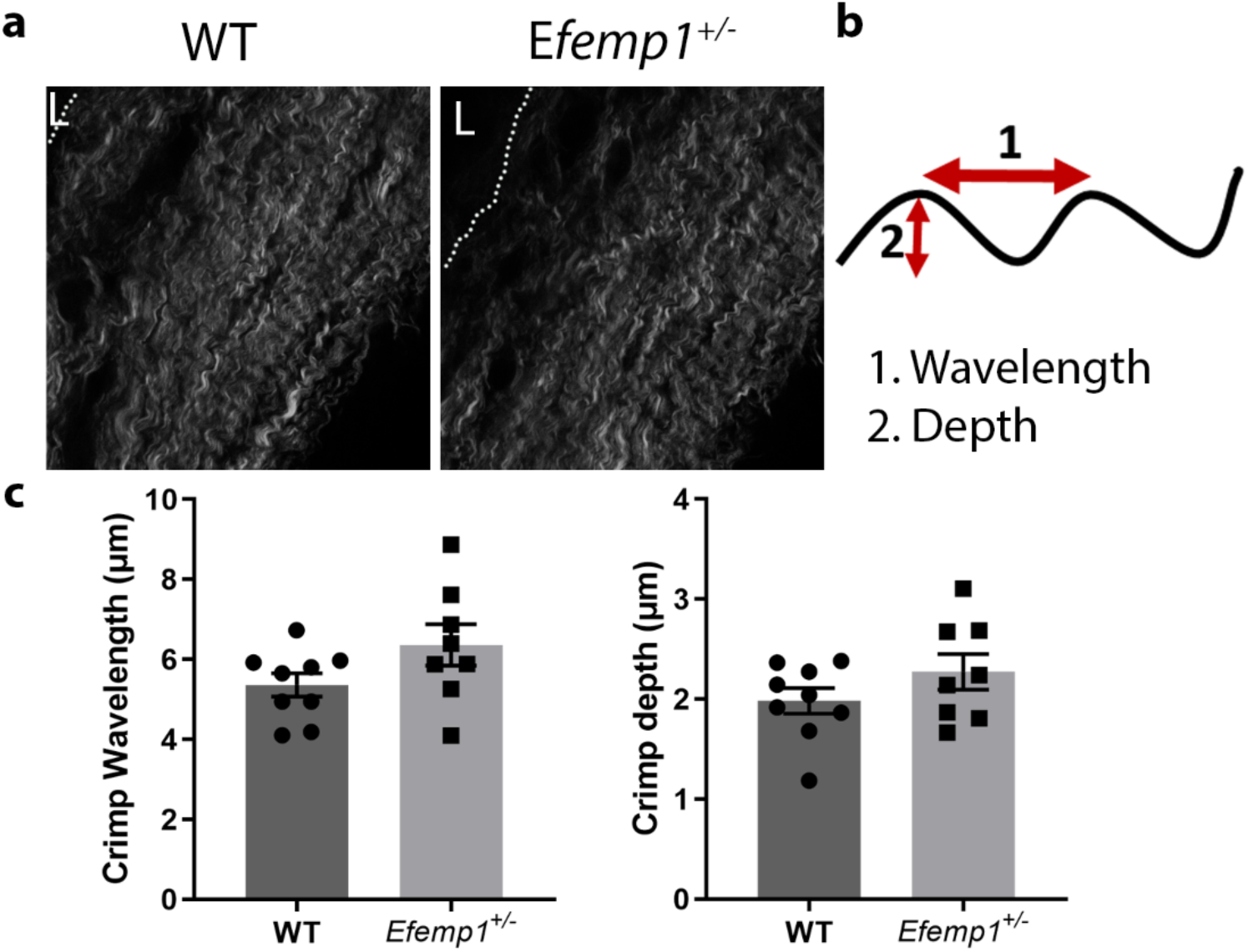
No difference in collagen fiber crimping between WT and *Efemp1^+/-^* extrahepatic bile ducts. (a) Representative images of SHG microscopy used for fiber analysis. Dotted lines and **L** denote lumens. (b) Schematic of fiber crimp wavelength and depth measurements used for analysis. (c) Quantification of crimping. There is no difference between WT and E*femp1^+/-^*crimp wavelength (p= 0.14041) or depth (p=0.1988). For each analysis, 15 crimps per image were analyzed in 3 images per duct. 8-9 individual ducts were examined per genotype. Data shown are mean ± SEM with each point representing the mean of a different duct. Significance was determined by unpaired T-test.

**Supplemental Figure 2:**
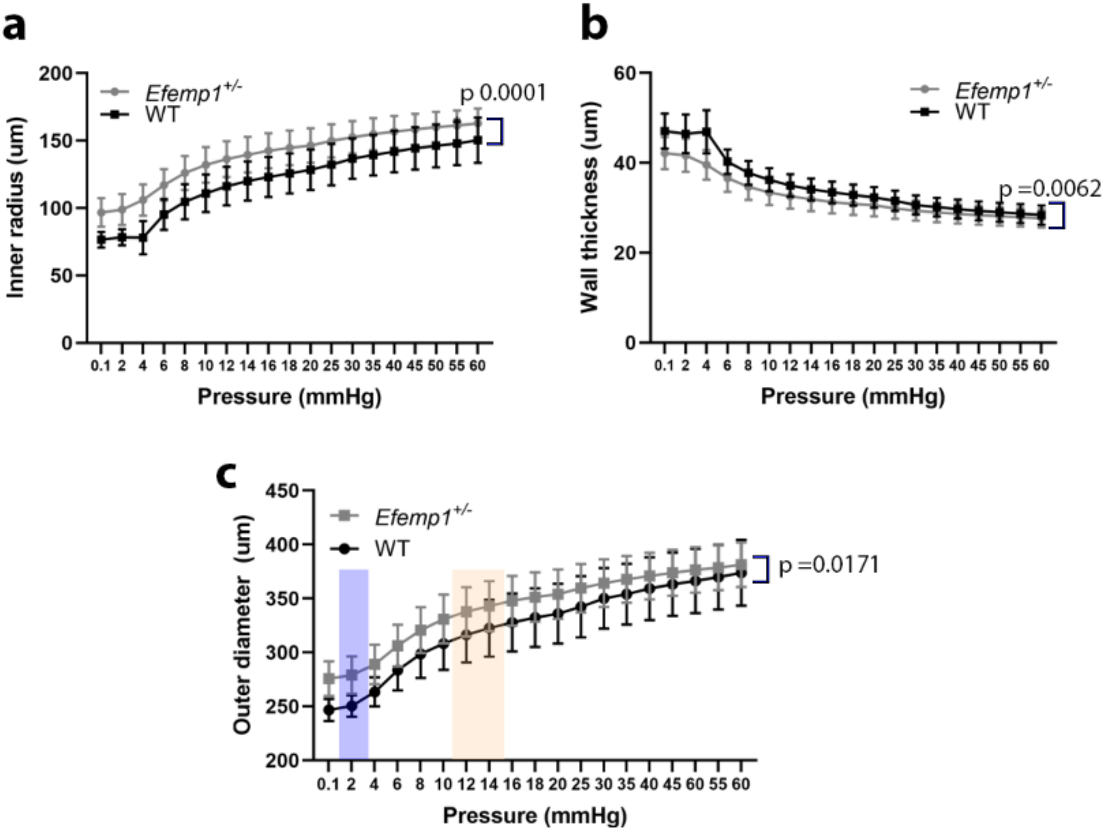
Altered circumferential mechanics of *Efemp1^+/-^* extrahepatic bile ducts. (a) Measured inner radius of extrahepatic bile ducts at increasing pressures. (b) Measured wall thickness of extrahepatic bile ducts at increasing pressures. (c) Measured outer diameter of extrahepatic bile ducts at increasing pressures. Pressure was increased in 2-mm increments from 0-20 mm Hg followed by 5-mm increments (from 20-60 mmHg). 6-7 individual extrahepatic bile ducts were tested per genotype. Data shown are mean ± SEM, two-way ANOVA with a Šidák post-hoc test statistical test.

